# Regeneration can take place across Drosophila compartments or segments with different Hox gene expression

**DOI:** 10.64898/2026.02.16.706018

**Authors:** Rafael Alejandro Juárez-Uribe, Paloma Martín, Laura Utiel, Blanca L. Arrabal, Marina Blanco, Roberto Yagüe-Serrano, Eduardo Cazalla, Ernesto Sánchez-Herrero

**Affiliations:** Department of Pathology and Laboratory Medicine, University of Rochester Medical Center, Rochester, New York, U.S.A.; Centro de Neurociencias Cajal (CNC), CSIC, Av. León 1, Alcalá de Henares, 28805 Madrid, Spain; Health Research Institute La Fe, Avenida Fernando Abril Martorell 106, 46026, Valencia, Spain; Department of Biochemistry, School of Medicine, Universidad Autónoma de Madrid (UAM), 28049 Madrid, Spain; Instituto de Investigaciones Biomédicas "Sols-Morreale" (CSIC-UAM), 28029 Madrid, Spain; Instituto de Investigación Sanitaria La Paz (IdiPaz), 28046 Madrid, Spain; Centro de Investigación Biomédica en Red de Enfermedades Neurodegenerativas (CIBERNED), 28046 Madrid, Spain

## Abstract

Regeneration in *Drosophila* has primarily been studied in imaginal discs, which are organized into distinct compartments defined by strict lineage boundaries. While cells typically do not cross these borders, regeneration can bridge them to restore the original organ specification, which is determined by Hox genes. Although differences in Hox gene expression are known to segregate cell populations, the role of such differences as a potential barrier to regeneration remains unclear. We investigated this by analyzing two experimental settings: the analia (derived from the Hox-compound genital disc) and haltere discs with mutations in *Ultrabithorax* regulatory regions (*bithorax* or *postbithorax*). Our findings demonstrate that Hox gene differences are not an absolute impediment to regeneration across segments or compartments. However, we observed occasional regenerative limits, which were non-specifically enhanced in the *postbithorax* background.

## Introduction

Regeneration is a process whereby an organism restores the structure and function of a part of the body that has been lost or damaged. Regeneration has been studied in different organisms, among them *Drosophila melanogaster*, in which the wing imaginal disc has been the main target of investigation (Smith-Bolton et al., 2009; Bergantiños et al., 2010; Worley et al., 2012).

One key question in regeneration studies is the origin of the cells that repair the damaged organ. Different experiments in *Drosophila* imaginal discs have demonstrated that cells from outside the injured domain contribute to its regeneration. For example, cells from the wing disc hinge contribute to the regenerated wing pouch (Smith-Bolton et al., 2009; Herrera et al., 2013; Verghese and Su, 2016). At a more local scale, cells from an intervein region of the wing pouch may be re-specified to replace vein cells that have undergone cell death (Repiso et al., 2013).

The wing disc, like other discs, is first divided into anterior (A) and posterior (P) compartments (García-Bellido et al., 1973). These are units of lineage separated from very early in development, so it was expected that regeneration in a disc would be carried out by cells from the same compartment that is injured. However, it was demonstrated that a damaged P compartment could be restored by cells coming from the A compartment, which changed compartment identity during this process (Herrera & Morata, 2014).

Specification of imaginal discs, and in general of different segments along the anteroposterior (A/P) axis, is provided by Hox genes (Lewis, 1978; Rezsohazy et al., 2015). The Hox gene Antennapedia is expressed in the wing disc (Paul et al., 2021), whereas *Ultrabithorax* (*Ubx*) differentiates the homologous haltere from the wing disc (Lewis, 1963). Several studies have indicated that Hox genes can induce cell segregation and separate cell populations (García-Bellido, 1968; Morata and García-Bellido, 1976; Estrada and Sánchez-Herrero, 2001; Gandille et al., 2010; Curt et al., 2006; Prin et al., 2014; Tiberghien et al., 2015). Therefore, it is unknown if regeneration across compartments, as described in the wing disc (Herrera and Morata, 2014), can also occur when the two compartments or segments have different homeotic information.

This question can be addressed if a mutation in a Hox gene results in compartment-specific Hox expression, or in discs, like the genital disc, which express more than one Hox gene. Previous studies in which the genital disc was mechanically fragmented proved its ability to regenerate (Ursprung, 1962; Bryant, 1975a; Bryant, 1975b; Littlefield and Bryant, 1979). However, these experiments could not delimit precisely the point of fragmentation, and therefore cannot ascertain the question of whether differences in Hox gene expression affect regeneration.

We have addressed this issue analyzing regeneration in adult derivatives of the genital disc and in haltere discs bearing *Ubx* regulatory mutations. Both cases are organs with compartments or segments with different Hox expression. Our results are consistent with transgression of segment boundaries during regeneration in adult male analia and in *bithorax* (*bx*) mutant haltere discs, but also show duplications of anterior haltere and wing discs tissue in a *postbithorax* (*pbx*) mutant background. This may be due to an unspecific increase in the frequency of such exceptional cases occurring in an *Ubx* wildtype background.

## Results

### The analia regenerates after cell death induced by expression of pro-apoptotic genes

The genital disc is made by the fusion of the A8, A9 and A10 abdominal segments (Nöthiger et al., 1977; Schübpach et al., 1978), which express the Hox gene *Abdominal-B* (*Abd-B*) in the A8 and A9 and *caudal* (*cad*) in the A10 (MacDonald and Struhl, 1986; Mlodzik and Gehring, 1987; Freeland and Kuhn, 1996; Casares et al., 1997; Estrada and Sánchez-Herrero, 2001; Morata and Moreno, 1999; Gorfinkiel et al., 2003; Foronda et al., 2006) (Fig. 1A-D’’). Although *cad* is not a Hox gene, it bears several features in common with Hox genes: it has a homeobox, it represses *Abd-B*, which is expressed just anterior to *cad* in the embryo and genital disc (following the Hox gene rule of transcriptional down-regulation along the A/P axis), and specifies a particular structure, the analia (Moreno and Morata, 1999). The restricted expression of *cad* and *Abd-B* to A10 and A9 (Fig. 1B’, B’’, D’, D’’), respectively, and their specification of analia and genitalia, prompted us to analyze if the regeneration of the analia may use cells coming from the A9 primordium.

**Figure 1.**
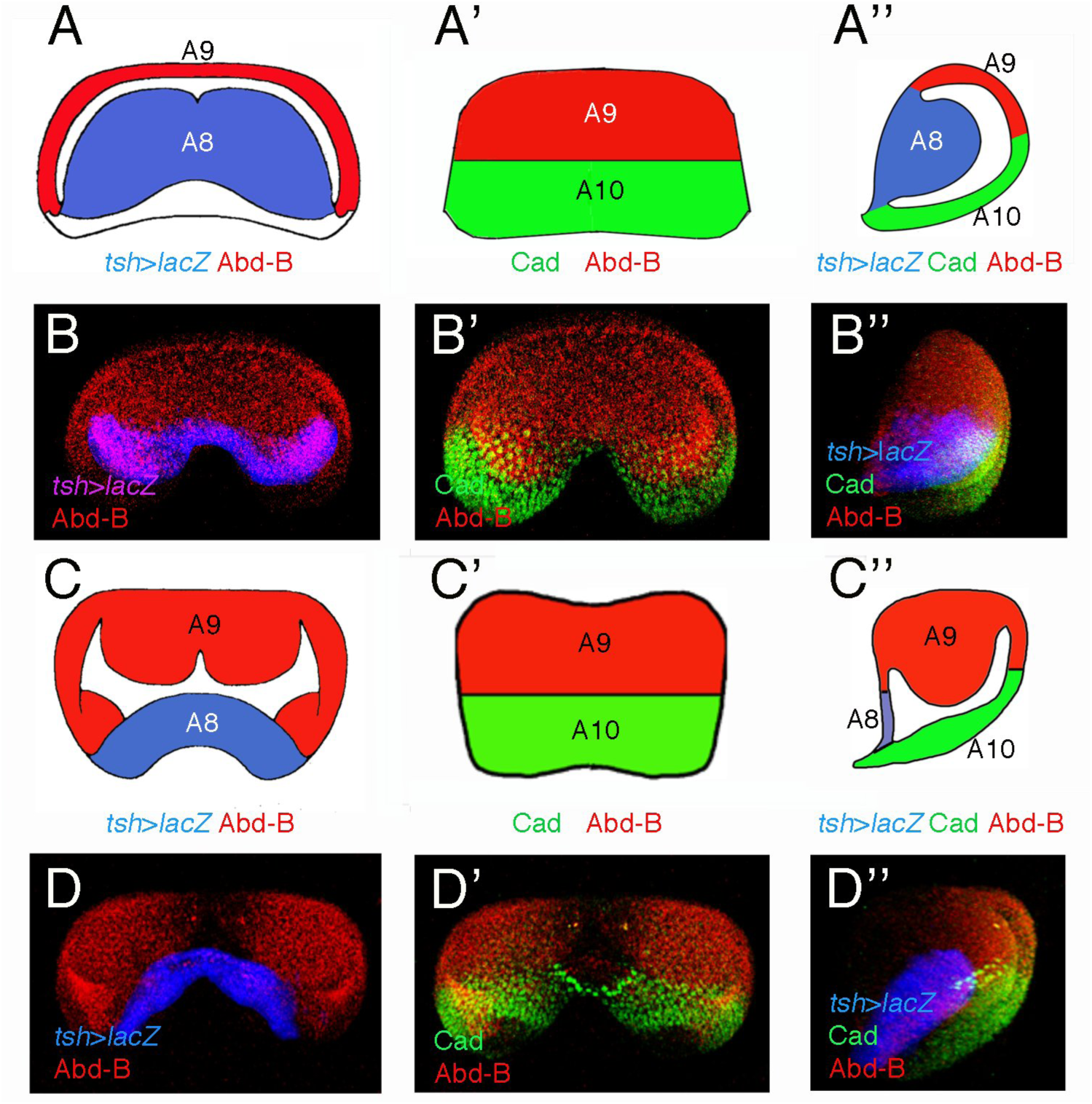
Segments in the genital disc. The female (A-B’’) and male (C-D’’) genital disc segments are represented with drawings (A-A’’, C-C’’) and their gene expression shown (B-B’’, D-D’’). The orientation of the discs is ventral (A-D), dorsal (A’-D’) and lateral (A’’-D’’). In the female disc the A8 is bigger than the A9 while in the male is the opposite, and in the A8 of both sexes there is expression of genes like *teashirt* (*tsh*) (Gorfinkiel et al., 2003). Abd-B is expressed in both A8 and A9 and Cad in A10.

We used the UAS/Gal4/Gal80^ts^ system (McGuire et al., 2003) to drive timing expression, under *cad*-Gal4 control, of the pro-apoptotic genes *head involution defective* (*hid*) or *reaper* (*rpr*), and studied ensuing regeneration (Smith-Bolton et al 2009; Bergantiños et al., 2010; Worley et al., 2012). Surprisingly, *cad*-Gal4 *tub*-Gal80^ts^ UAS-*hid* or *cad*-Gal4 *tub*-Gal80^ts^ UAS-*rpr* females showed analia with very thin and near transparent bristles when grown at 18°C (Fig. 2C, D, compare with the wildtype and *cad*-Gal4/+ females grown at the same temperature, Fig. 2A, B). Males, by contrast, showed the same phenotype as the controls (Fig. 2E-H). Therefore, the effect of cell death on bristle development cannot be completely suppressed by *tub*-Gal80ts at 18°C, as is the case in other body parts (Supplementary Figure 1). Animals in which cell death was induced at the third larval period lack analia (Fig. 2I, K, M, O), or both genitalia and analia if induction was done earlier in development (Fig. 2J, L, N, P). This effect on genitalia is probably due to the interaction between the genitalia and analia primordia (Gorfinkiel et al 2003), and is more frequent in males, perhaps because those primordia are contiguous in the male genital disc.

**Figure 2.**
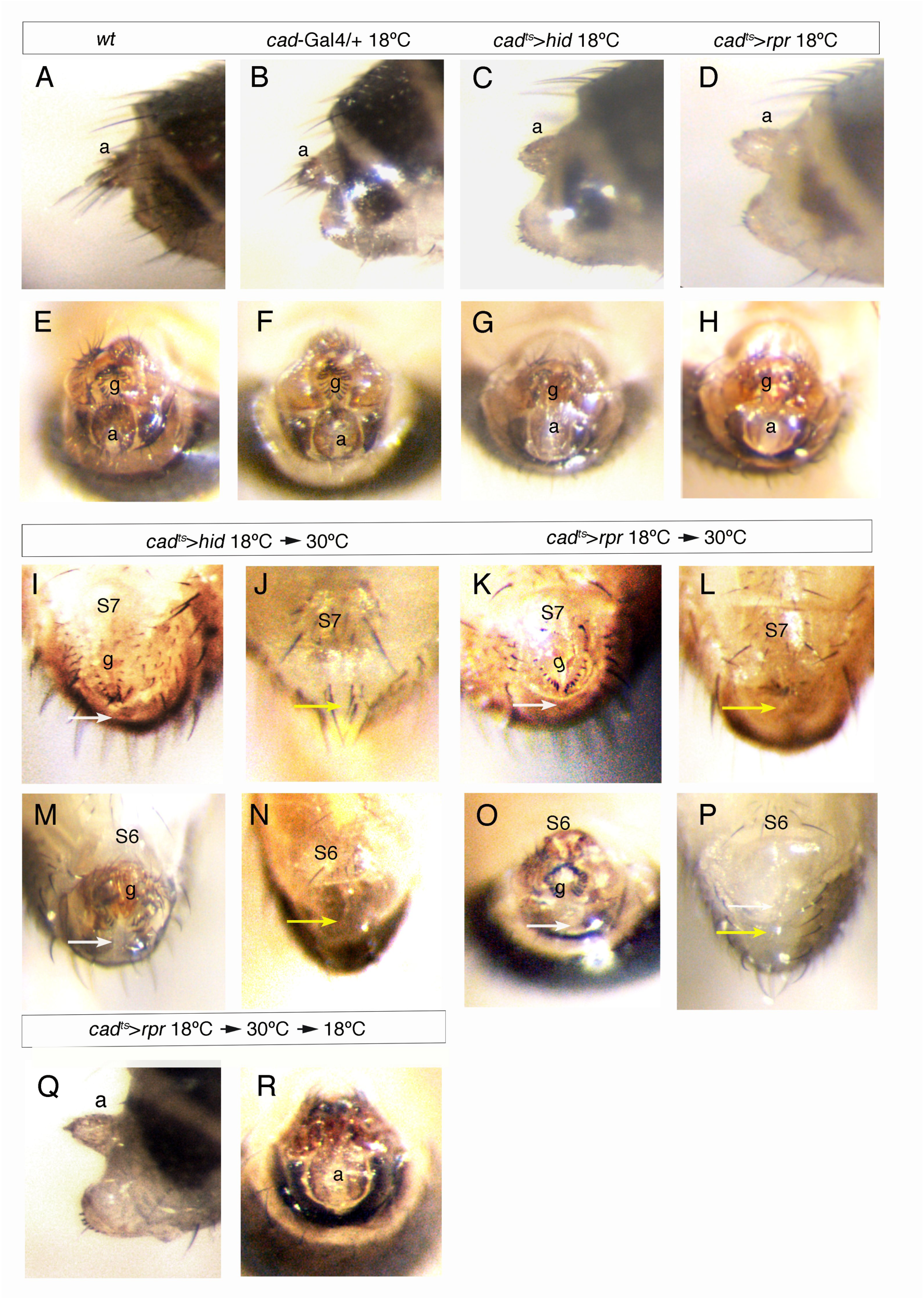
Development of genitalia and analia in males and females after cell death induced in the analia. (A-H) Lateral (A-D) and ventral (E-H) views of the posterior abdominal segments and analia (a) of females (A-D), and genitalia (g) and analia of males (E-H), of *wt* (A, E), *cad*-Gal4/+ at 18°C (B, F), *cad*-Gal4/UAS-*hid tub*-Gal80^ts^ at 18°C (C, G) and *cad*-Gal4 *tub*-Gal80^ts^/UAS-*rpr* at 18°C (D, H). (I-P) Ventral views of the posterior part of females (I-L) and males (M-P) of *cad*-Gal4/ UAS-*hid tub*-Gal80^ts^ (I, J, M, N) and *cad*-Gal4 *tub*-Gal80^ts^/UAS-*rpr* (K, L, O, P) shifted from 18°C to 30°C at early or late third instar. (Q, R) Female (Q) and male (R) *cad*-Gal4 *tub*-Gal80^ts^/UAS-*rpr* analia after cell death induction and regeneration, showing the “wt” phenotype in females and normal analia in males. White arrows indicate absence of analia and the yellow arrows, of genitalia and analia; g, genitalia. S6 and S7, sixth and seventh sternites, respectively. The >symbol in this and subsequent Figures represents the Gal4/UAS combination.

After the cell death and regeneration protocol (see Methods) females could recover the analia observed when grown at 18°C (what we call “wt”, see Fig. 2C, D, Q), whereas males can fully recover the wildtype analia (Fig. 2R). We also note, when inducing apoptosis with *hid*, a better regeneration in females (reaching the “wt” phenotype) than in the males (see Table 1).

**Table 1.** Regeneration numbers after cell death induced in the caudal domain expressing the proapoptotic gene *hid*. The cross is *cad*-Gal4/*CyO, Tb* x UAS-*hid tub*-Gal80^ts^/*CyO, Tb*. Treatments are 24h at 18°C, 5 or 6 days (d) at 18°C and 30h or 40h at 30°C before shifting back the larvae to 18°C. “wt” indicates females with the analia phenotype of females at 18°C with *tub*-Gal80^ts^ (see text). No A/No A + G means absence of analia or of genitalia and analia, respectively.

| Treatment |  | hid | hid |
| --- | --- | --- | --- |
|  |  | females | males |
| 24h-6d-30h | "wt" or wt | 423 | 203 |
|  | partial | 26 | 16 |
|  | No A/No A+G | 96 | 213 |
|  | "wt" or wt/total ratio | 423/545<br>0,77 | 144/432<br>0,33 |
| 24h-5d-40h | "wt" or wt | 68 | 31 |
|  | partial | 10 | 10 |
|  | No A/No A+G | 14 | 72 |
|  | "wt" or wt/total ratio | 68/92<br>0,73 | 31/113<br>0,27 |

We examined to which extent the treatment used was really producing cell death and regeneration in the larval A10 primordium. We checked apoptosis with an anti-cleaved Dcp-1 antibody and trace the lineage of cells expressing *cad* with the flip-out technique (Struhl and Basler, 1993) in larvae of the genotype *cad*-Gal4/UAS-*hid tub*-Gal80^ts^; UAS-*flp act*>stop>*lacZ*/+. At 18°C, since the Gal80^ts^ protein inhibits Gal4 activity, no high levels of cell death or marked cells were observed (Supplementary Fig. 2A-A’’), whereas after a 24h pulse at 30°C the abnormal genital discs show massive expression of activated Dcp-1, roughly coincident with *cad*-Gal4 cell lineage (Supplementary Fig. 2B-B”). The activation of the c-Jun N-terminal kinase (JNK) pathway (Bergantiños et al., 2010; Herrera et al., 2013), revealed with the TRE-DsRed reporter (Chatterjee and Bohmann, 2012), and that of the Wg activity BRV-B-GFP reporter (Harris et al., 2016), are detected, in controls, in a small region (stalk) connecting the genital disc with larval tissue (Macías et al., 2004; Rousset et al., 2010), but extension into the A10 segment is observed when cell death is induced (Supplementary Fig. 2C-C”, D-D’’).

### Cells outside the *cad* domain probably contribute to the regeneration of the analia

To know if cells cross the A9/A10 boundary during regeneration of the analia primordium, we first observed Abd-B and Cad expression in the male and female genital discs. As shown in Fig 3A-A’’’, B-B’’’, there is a slight overlap between Abd-B-expressing and Cad-expressing cells at the A9/A10 boundary, both in male and female genital discs. This overlap has been noticed before (Moreno and Morata, 1999), and makes it difficult to know if cells from one primordium contribute to the regeneration of the other. Moreover, the *cad*-Gal4 line does not precise coincide with Cad (Fig. 3C-C’’’), with a few Cad-expressing cells being located more anteriorly than those in which the *cad*-Gal4 line is active. These drawbacks hampers detecting if cells from the A9 segment, expressing *Abd-B*, change segment position and Hox expression to regenerate the A10 segment.

**Figure 3.**
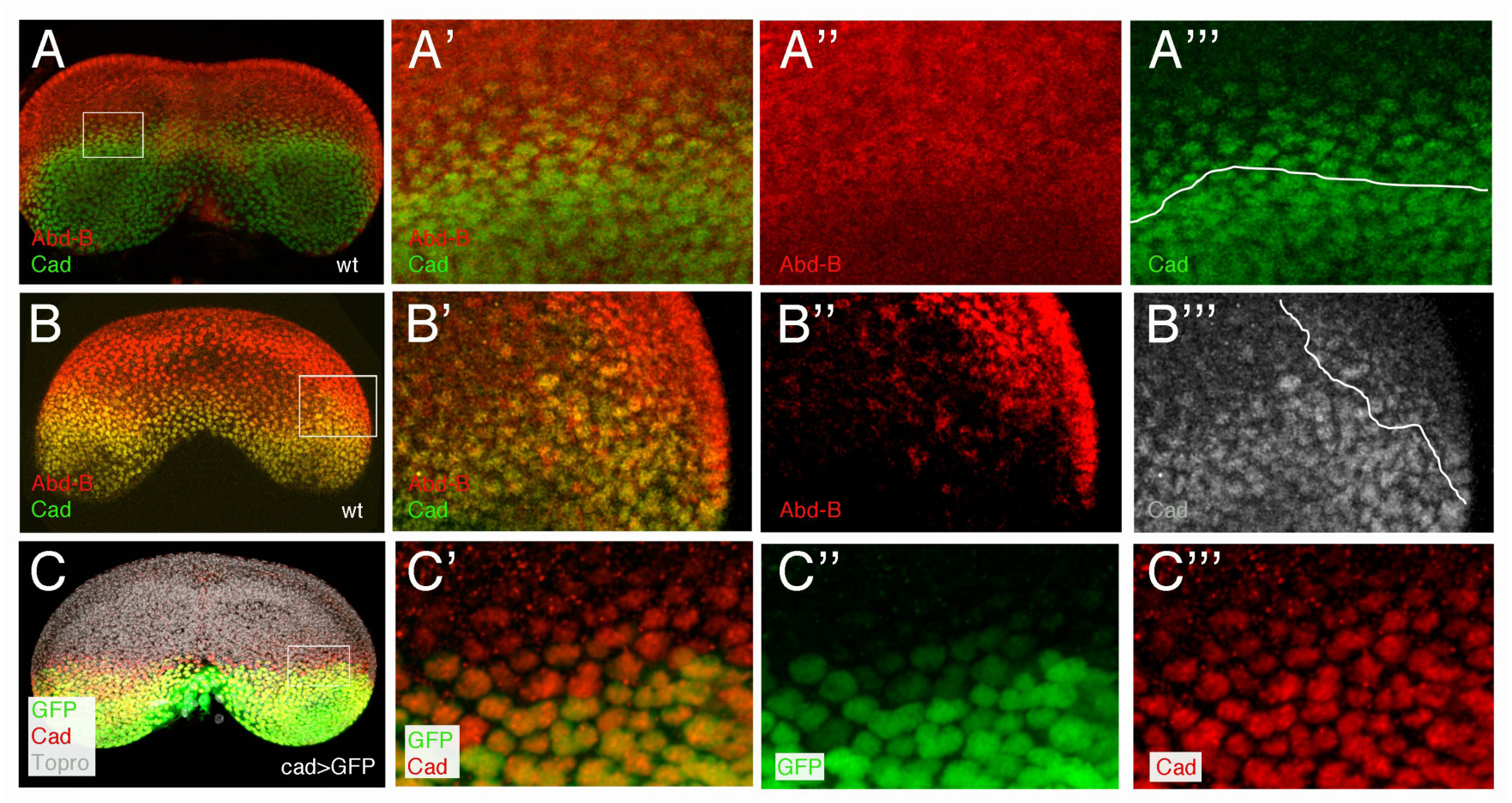
Caudal and Abdominal-B expression overlap in the larval genital discs. (A-B’’’) Abd-B (red) and Cad (green) expression in the third instar male (A-A’’’) and female (B-B’’’) genital discs. The insets in A and B are amplified in A’-A’’’ and B’-B’’’, respectively, and show the overlap in the expression of the two proteins. (C-C’’’) *cad*-Gal4 UAS-*GFP* third instar female disc stained with anti-Cad, showing Cad expression anterior to GFP expression (the inset in C is amplified in C’-C’’’).

We turned then to the study of the adult structure. Contrary to what is observed in third instar genital discs, in pupa the expression of Abd-B and Cad is non-overlapping (McQueen et al., 2025). Clones induced early in development are confined to either genitalia or analia, indicating early lineage segregation between the two structures (Dübendorfer and Nöthiger, 1982). Consistently, the *cad*-Gal4 line direct *yellow* expression in the analia and not in the adjacent genital structures, both in males and females (Fig. 4A, B).

**Figure 4.**
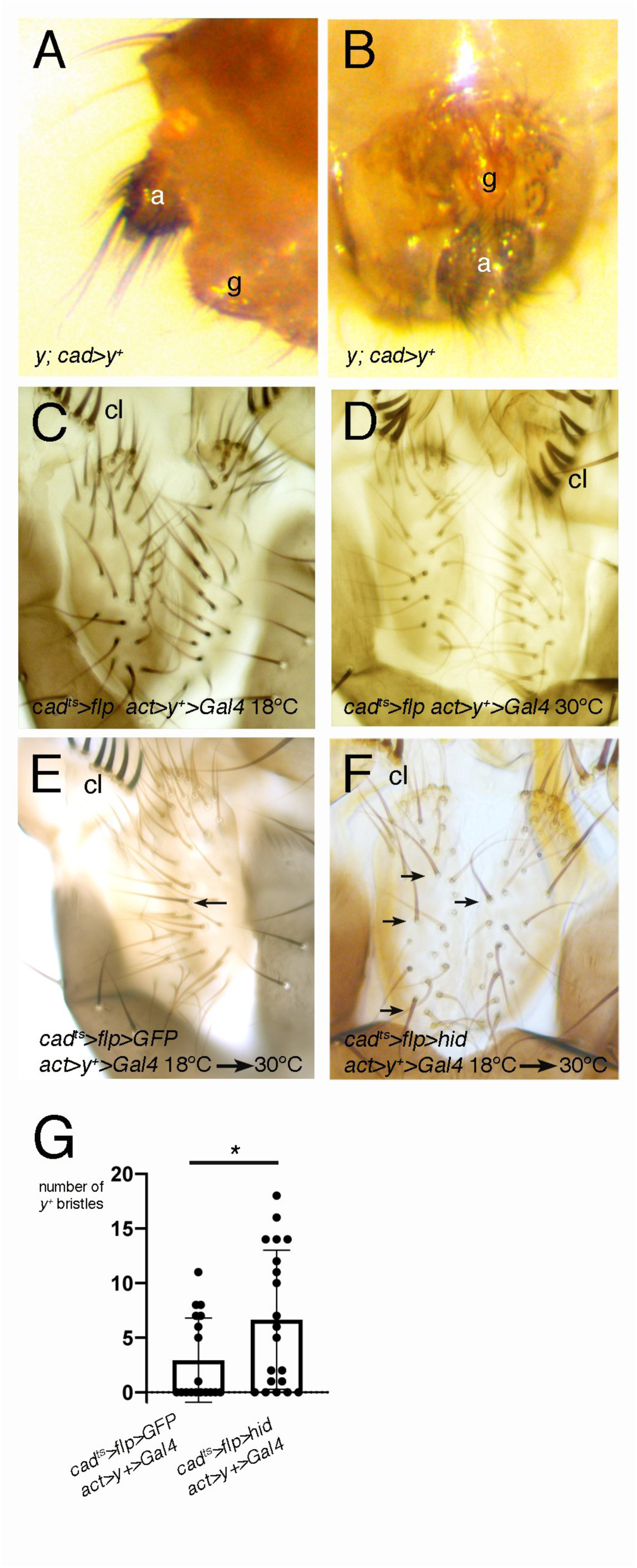
Increase of *yellow* mosaicism in the analia after cell death and regeneration. (A, B) Analia of females (A) and males (B) in adults of the genotype *y*; *cad*-Gal4/UAS-*y^+^*, showing the restriction of the dark tissue (*y^+^*) to the analia (a); g, genitalia. (C) Analia of a male of the genotype *y w*; *cad*-Gal4 *tub*-Gal80^ts^/ *act*>*y+*>Gal4; UAS-*flp*/*+* grown at 18°C, showing the bristles are of brown color (no recombination of the cassette that liberates the *y^+^* gene (n=6). The FRT sequences are represented by the symbol > (which also indicates Gal4/UAS). In this and following panels cl means clasper teeth. (D) Analia of a male of the same genotype as in C but grown at 30°C: in this case the Gal4 activates the flipase in the analia, and this leads to recombination of the cassette and losing of the *y^+^* gene, so that the whole analia is of yellow color (n=6). (E) Only one *y^+^* bristle (arrow) is observed in a male analia of the control genotype, *cad*-Gal4/*tub*-Gal80^ts^ UAS-*GFP*; UAS-*flp*/actin>*y^+^*>Gal4 (n=20), after the regeneration treatment. (F) In the analia of males of the genotype *cad*-Gal4/UAS-*hid tub*-Gal80^ts^; UAS-*flp*/actin>*y^+^*>Gal4, several *y^+^* bristles are observed (arrows) (n=18) with the same regeneration treatment.

To analyze if cells change segment identity during regeneration, we induced cell death and regeneration while marking the caudal lineage by the loss of the *yellow* (*y*) cuticular marker, present in the cassette *actin*-FRT-*y^+^*-FRT-Gal4 (abbreviated as *act*>*y+*>Gal4), in a *y* mutant background. Adults of the genotype *y w*; *cad*-Gal4 *tub*-Gal80^ts^/*act*>*y+*>Gal4; UAS-*flp*/*+* kept at 18°C do not have any *y* bristle in the analia (Fig. 4C), but this organ is full of *y* bristles when raised at 30°C (Fig. 4D). Following the regeneration protocol, in 55,5% of control males (*cad*-Gal4/*tub*-Gal80^ts^ UAS-*GFP*; UAS-*flp*/*act*>*y+*>Gal4) (n=18) the analia are completely *y*, while the rest of males have a small number of *y^+^* bristles, with an average of 2,9 *y^+^* bristles per analia (Fig. 4E, G). By contrast, in the experimental males (*cad*-Gal4/*tub*-Gal80^ts^ UAS-*hid*; UAS-*flp*/*act*>*y+*>Gal4) (n=20), only 25% of them have all the analia bristles *y*, and the average number of *y^+^* bristles is significantly higher, 6,6 (Fig. 4E, G). Although in some of the analia cells there might have been no removal of the cassette (in both groups), the data suggest that many *y+* bristles in the experimental set are probably derived from cells in the adjacent genitalia that expressed *Abd-B* before regeneration, and lose this expression to gain *cad* and develop analia.

### Cells can gain *Ultrabithorax* expression in regenerating *bithorax* haltere discs

To extend our results, we examined regeneration in haltere discs with compartment-specific Hox expression. The haltere disc is homologous to the wing disc but expresses *Ubx* (White and Wilcox, 1984; 1985; Cabrera et al., 1985; Beachy et al., 1985). In larvae mutant for the *bithorax* (*bx*) or *postbithorax* (*pbx*) *Ubx* regulatory regions, the anterior or posterior compartment of the disc, respectively, strongly reduces *Ubx* expression and is transformed into the corresponding one of the wing disc (Lewis, 1963; Cabrera et al., 1985; White and Wilcox, 1985). Therefore, the A/P boundary separates compartments of different fate, haltere or wing, with or without *Ubx* expression. We investigated whether, during regeneration of the posterior compartment of *bx* or *pbx* haltere discs, cells from the anterior compartment could gain (in *bx*) or lose (in *pbx*) *Ubx* expression as they rebuild the damaged compartment. The *hh*-Gal4 line was used as a posterior specific Gal4 driver for these experiments.

Before analyzing results in *bx* or *pbx* mutants, we examined lineage tracing in wildtype wing and haltere discs. In larvae of the genotype UAS-*y^+^*/*+*; UAS-*flp act*>*stop*>*lacZ*/*hh*-Gal4 *tub*-Gal80^ts^, shifted during second larval instar to 30°C for 30h, we confirmed that cells marking the posterior lineage abut the anterior ones, marked by Cubitus interruptus (Ci) (Eaton and Kornberg, 1990), in both wing and haltere Fig. S3A-B’’’) (Table 2).

**Table 2.** Number of wing (w) and haltere (h) discs undergoing regeneration, showing partial duplications of anterior compartment, or abnormal development in the different genotypes indicated. *Two of the haltere discs show crossing of anterior cells to the posterior compartment; **In 12 discs there are a few posterior cells in the anterior compartment. ***There are cases of reduced posterior compartment and abnormal compartment boundary (more easily observed in the wing disc).

| Genotype | Total | Normal regeneration | Tissue duplications | Abnormal development |
| --- | --- | --- | --- | --- |
| UAS-*y<sup>+</sup>*/+; UAS-*flp*<br>*act>stop>lacZ*/hh-Gal4 *tub*-Gal80<sup>ts</sup> -w | 24 | 24 | 0 | 0 |
| UAS-*y<sup>+</sup>*/+; UAS-*flp*<br>*act>stop>lacZ*/hh-Gal4 *tub*-Gal80<sup>ts</sup> -h | 21 | 21 | 0 | 0 |
| *ubi>stop>GFP* UAS-*flp*/UAS-*lacZ*<br>*tub*-Gal80<sup>ts</sup>; *bx<sup>3</sup>* hh-Gal4/TM2- h | 11 | 11 | 0 | 0 |
| *ubi>stop>GFP* UAS-*flp*/UAS-*hid*<br>*tub*-Gal80<sup>ts</sup>; *bx<sup>3</sup>* hh-Gal4/TM2- h | 18 | 18* | 0 | 0 |
| *ubi>stop>GFP* UAS-*flp*/UAS-*lacZ*<br>*tub*-Gal80<sup>ts</sup>; *pbx<sup>1</sup>* hh-Gal4/TM2- h | 25 | 25** | 0 | 0 |
| *ubi>stop>GFP* UAS-*flp*/UAS-*hid*<br>*tub*-Gal80<sup>ts</sup>; *pbx<sup>1</sup>* hh-Gal4/TM2 - h | 21 | 13 | 8 | 0 |
| UAS-*y<sup>+</sup>*/UAS-*flp* *act>stop>lacZ*; *ci*-Gal4 *tub*-Gal80<sup>ts</sup> -w | 15 | 15 | 0 | 0 |
| UAS-*hid* *tub*-Gal80<sup>ts</sup>/ UAS-*flp*<br>*act>stop>lacZ*; *ci*-Gal4/+ -h | 18 | 18 | 0 | 0 |
| UAS-*hid* *tub*-Gal80<sup>ts</sup>/+; *ci*-Gal4<br>*bx<sup>3</sup>*/TM2 -h | 22 | 21 | 1 | 0 |
| UAS-*hid* *tub*-Gal80<sup>ts</sup>/+; hh-Gal4<br>*pbx<sup>1</sup>*/TM2-w | 27 | 8*** | 14 | 5 |
| UAS-*hid* *tub*-Gal80<sup>ts</sup>/+; hh-Gal4<br>*pbx<sup>1</sup>*/TM2-h | 14 | 7*** | 4 | 3 |
| UAS-*hid* *tub*-Gal80<sup>ts</sup>/; hh-Gal4/<br>UAS-*flp* *act> stop>lacZ* -w | 18 | 17*** | 1 | 0 |
| UAS-*hid* *tub*-Gal80<sup>ts</sup>/; hh-Gal4/<br>UAS-*flp* *act> stop>lacZ* -h | 15 | 15*** | 0 | 0 |

Then, we investigated regeneration in haltere discs mutant for *bx^3^* (*bx^3^*/*TM2*), which show a strong reduction of *Ubx* in the anterior compartment (Fig. S4A, A’). Apoptosis induction was confirmed by massive Dcp-1 signal in the P compartment, coinciding with the *hh* lineage (Fig. S4C, C’). We next analyzed possible changes in *Ubx* expression after regeneration of these discs. In the control discs (expressing *lacZ* instead of *hid*), the *hh* lineage coincides with the *Ubx* expression in the posterior compartment (Fig. 5A-A’’’) (Table 2). In most of the experimental larvae (16/18), we observed regeneration of the posterior compartment only with cells expressing both *Ubx* and the *hh* lineage. However, in 2 haltere discs we noted a group of cells apparently integrated in the posterior compartment and close to the A/P boundary that express *Ubx* but lack GFP signal, that is, they are not part of the *hh* lineage (Fig. 5B-B’’) (Table 2). These cells are likely to be anterior cells that penetrate into the posterior compartment and acquire *Ubx* expression.

**Figure 5.**
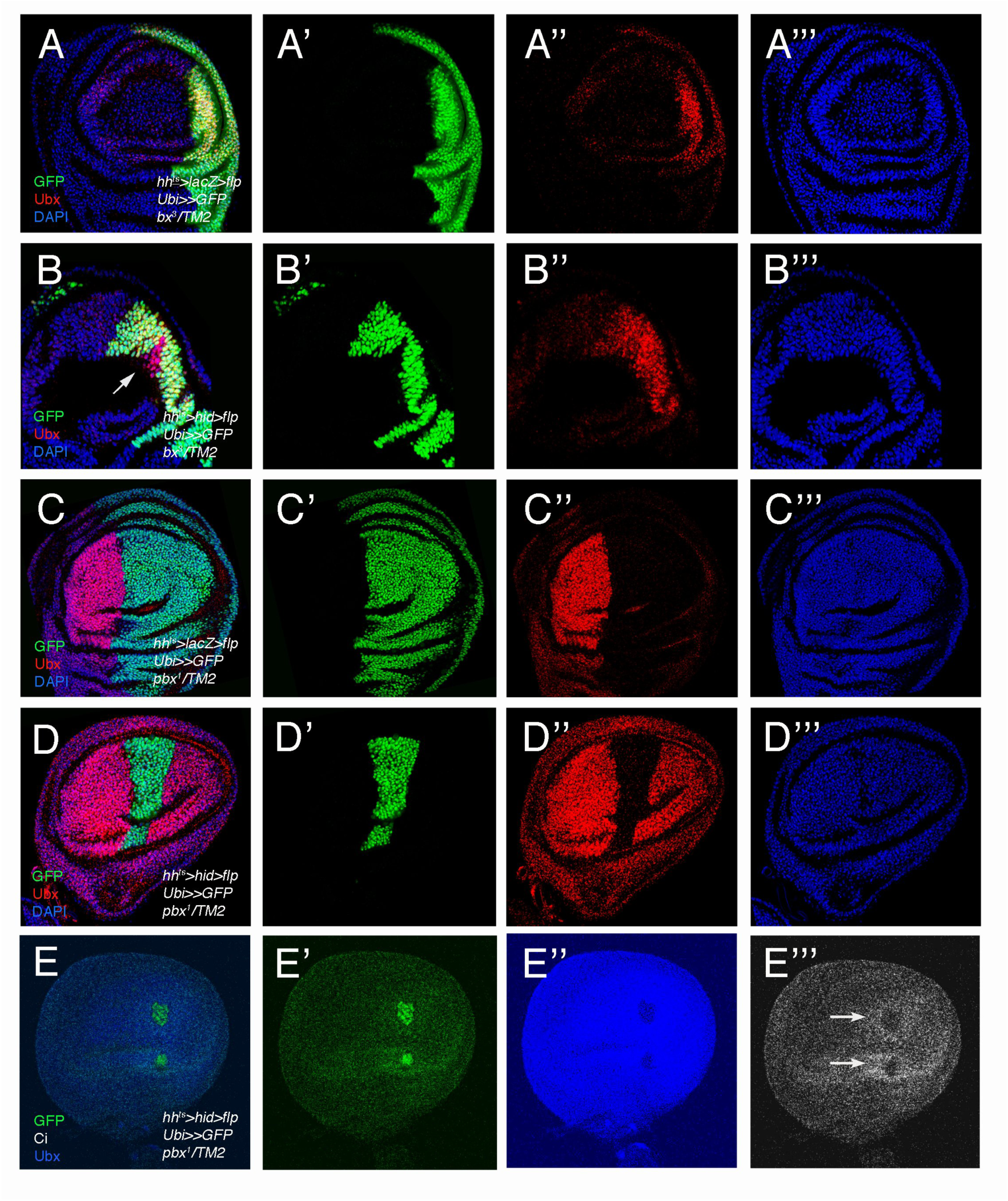
Regeneration and duplication of haltere disc tissue after cell death and recovery in the posterior compartment. (A-A’’’) Third instar haltere disc mutant for *bx* in which *lacZ* is expressed in the posterior compartment and the cell lineage in the same compartment is marked by GFP expression (genotype *ubi*>stop>*GFP* UAS-*flp*/UAS-*lacZ tub*-Gal80^ts^; *bx^3^ hh*-Gal4/*TM2*). Ubx is almost completely absent from the anterior compartment and its expression coincides with that of GFP. In this and following Figures the double arrowhead (>>) stands for FRT-stop-FRT. (B-B’’’) Third instar haltere disc of the genotype *ubi*>stop>*GFP* UAS-*flp*/UAS-*hid tub*-Gal80^ts^; *bx^3^ hh*-Gal4/*TM2* showing some anterior cells (not marked with the GFP lineage tracer) integrated in the posterior compartment and showing Ubx antibody staining (arrow). (C-C’’’) Third instar haltere disc mutant for *postbithorax* in which *lacZ* is expressed in the posterior compartment and the cell lineage in the same compartment is marked by GFP expression (genotype *ubi*>stop>*GFP* UAS-*flp*/UAS-*lacZ tub*-Gal80^ts^; *pbx^1^ hh*-Gal4/*TM2*). Ubx antibody signal is observed just in the anterior compartment, complementary to that of GFP. (D-E’’’). Two examples of haltere discs in which duplications are observed. The genotype of the discs is *ubi*>stop>*GFP* UAS-*flp*/UAS-*hid tub*-Gal80^ts^; *pbx^1^ hh*-Gal4/*TM2*. The cell lineage of the posterior compartment is marked by GFP, and is surrounded by anterior cells marked by Ubx (D’’, E’’) or Ci (E’’’). The reduction of the posterior compartment is extreme in the haltere disc in E-E’’’.

### Duplications in haltere discs after cell death and recovery in one compartment

In the complementary experiment, in which we used the same experimental setup but in haltere discs of *pbx^1^*/*TM2* larvae, without *Ubx* in the posterior compartment (Fig. S4B, B’), we obtained a different result. With the same experimental procedure as in *bx* discs, we also observed strong anti-Dcp-1 signal in the posterior compartment, which coincides with the *hh* lineage (Fig. S4D, D’). In most control discs, the absence of *Ubx* in the posterior compartment strictly coincides with the *hh* lineage (Fig. 5C-C’’’), but in near half of these discs (48%) we note the GFP signal in a few cells near the A/P boundary that are also expressing *Ubx* (Fig. S5A-A’’’) (Table 2). This can perhaps be explained by the occasional early expression of the *hh*-Gal4 driver in the anterior compartment in this mutant combination, similar to what has been observed in the wildtype wing disc (Bosch et al., 2016).

In the experimental discs (genotype *ubi*>stop>*GFP* UAS-*flp*/UAS-*hid tub*-Gal80^ts^; *pbx^1^ hh*-Gal4/*TM2*) most of the discs (65%) regenerate the posterior compartment with cells not expressing *Ubx*, as in controls. However, in 38% of the discs we observed a quite different result: these discs show different degrees of reduction of the posterior compartment, confined to the most central region of the discs, with the most posterior cells lacking the *hh* lineage and expressing *Ubx*; that is, the posterior compartment seems to be surrounded by anterior cells (Fig. 5D-D’’’). The haltere disc, therefore, seems to adopt a sort of A-P-A disposition (what we call compartment “duplications”) (Table 2). The anterior specification of the more posterior cells in these discs was confirmed by staining with anti-Ci antibody, which marks in the wildtype the anterior compartment, with higher levels in cells abutting the A/P boundary (Motzny and Holmgreen, 1995; Aza-Blanc et al., 1997). This increased signal is limiting or surrounding posterior cells, which in some cases are reduced to just small groups (Fig. 5E-E’’’; Fig. S5B-B’’’). Collectively, the data suggest that the discs are forming a partial mirror image duplication of the anterior compartment.

As the formation of these duplications was unexpected, we tested if a similar effect took place in a mutant anterior compartment that also suffers cell death induction. In control (*ci*-Gal4 *tub*-Gal80^ts^ UAS-*y^+^* UAS-*flp act*>*stop*>*lacZ*) or experimental (*ci*-Gal4 *tub*-Gal80^ts^ UAS-*hid* UAS-*flp act*>stop>*lacZ*) larvae which undergo the same experimental treatment as the *hh*-Gal4 larvae, we observed the coincidence of Ci and ß-galactosidase (the anterior cell lineage) expression in haltere discs (Fig. 6A-A’’’; 6B-B’’’) (Table 2). We next induced cell death and recovery but in a *bx* mutant background. Although in these experiments we did not label the anterior cell lineage, by staining with anti-Ubx and anti-Ci antibodies we observed that, in all but one disc, there was the expected regeneration of the anterior compartment (Fig. 6C-C’’’). In this single case, however, we found a mirror image duplication (with *Ubx* signal at the lateral sides of the disc), being now P-A-P, similar to some of those we have described for the *pbx* haltere discs after cell death induction in the posterior compartment (Fig. 6D-D’’’) (Table 2). Taken together, the experiments suggest there are cases, particularly when inducing cell death in the posterior compartment of *pbx* haltere discs, in which there is little or almost no regeneration. Instead, the ablated compartment is reduced, becoming surrounded by the cells of the adjacent one.

**Figure 6.**
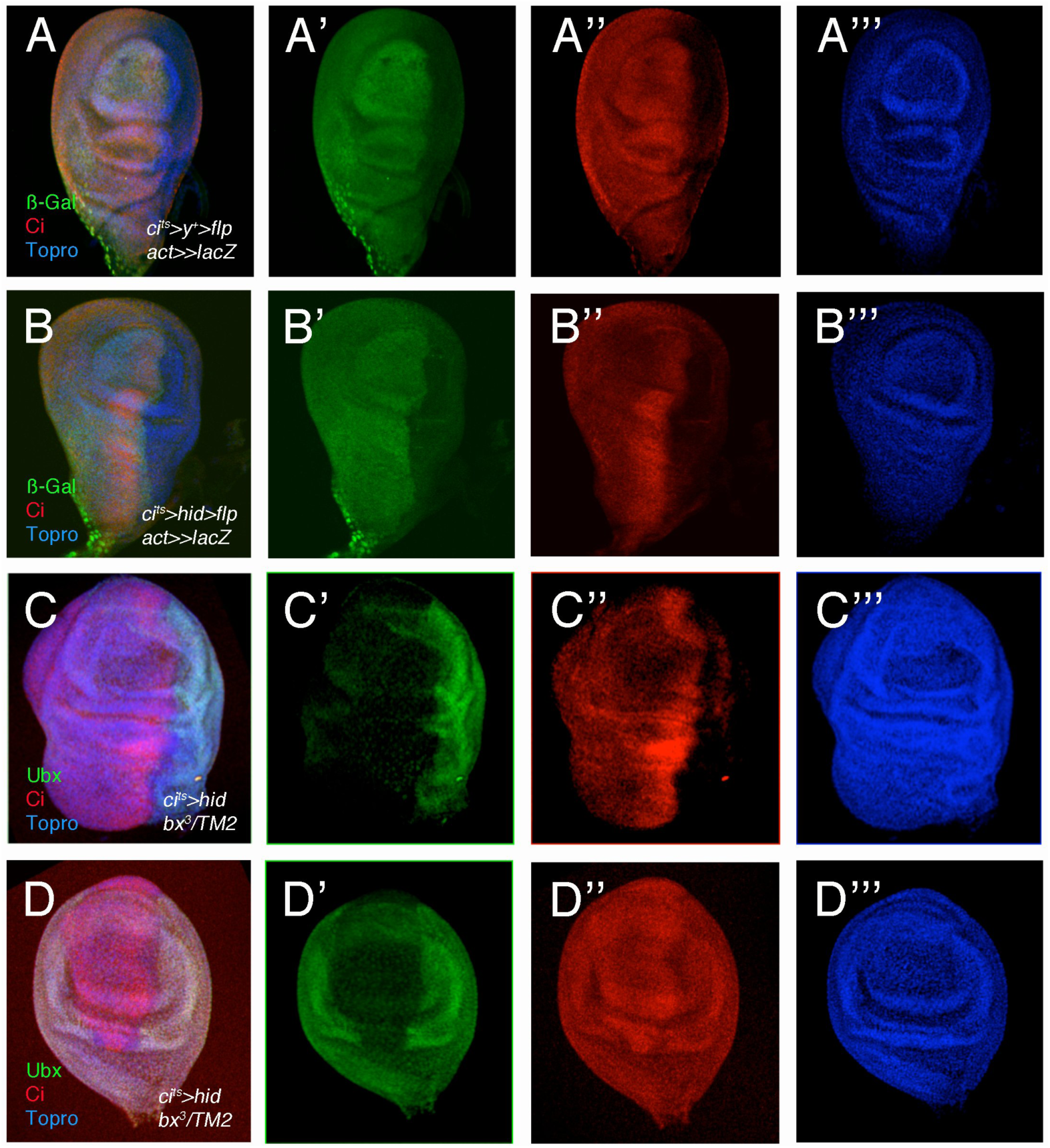
Development of *bithorax* mutant haltere discs after cell death and recovery in the anterior compartment. (A-A’’’) Control haltere discs of genotype *tub*-Gal80^ts^/UAS-*y^+^*; *ci*-Gal4/UAS-*flp act*>stop>*lacZ*. The anti-ß-galactosidase staining in the anterior compartment, marking the cell lineage, coincides with Ci signal (n=15). (B-B’’) The coincidence of cell lineage and Ci staining (UAS-*hid tub*-Gal80^ts^/+; *ci*-Gal4/UAS-*flp act*>stop>*lacZ)* is also observed in haltere discs in which there is cell death and recovery in the anterior compartment (n=18). (C-C’’’) Haltere disc mutant for *bx* in which cell death and recovery were induced in the anterior compartment (genotype UAS-*hid tub*-Gal80^ts^/+; *ci*-Gal4 *bx^3^*/*TM2*. There is normal Ubx and Ci expression. This occurs in 21/22 discs observed. (D-D’’’) Haltere disc of the same genotype as that in C but which shows duplication in the disc: posterior cells, marked by Ubx staining, surround anterior cells expressing Ci (there is high background in the Ci staining, but high levels at the A/P interphase could still be distinguished).

### Lack of complete regeneration after cell death in one compartment of the wing and haltere discs

To add new experimental evidence, we induced regeneration in larvae of the genotype UAS-*hid tub*-Gal80^ts^/+; *pbx^1^ hh*-Gal4/*TM2*, in which we might observe the presence of “duplications” even though we could not follow the cell lineage. Among the 14 *pbx* haltere discs analyzed, we observed 4 with “duplications” (Fig. 7A-A’’) and 3 have abnormal morphologies (Fig. S6A-A’’’). Surprisingly, we also found similar effects in the wing disc. Out of 27 discs analyzed, we found 14 with some kind of reduction of the posterior compartment and “duplication” of the anterior domain (Fig. 7B-B’’) and 5 with abnormal development (Fig. S6B-B’’) (Table 2).

**Figure 7.**
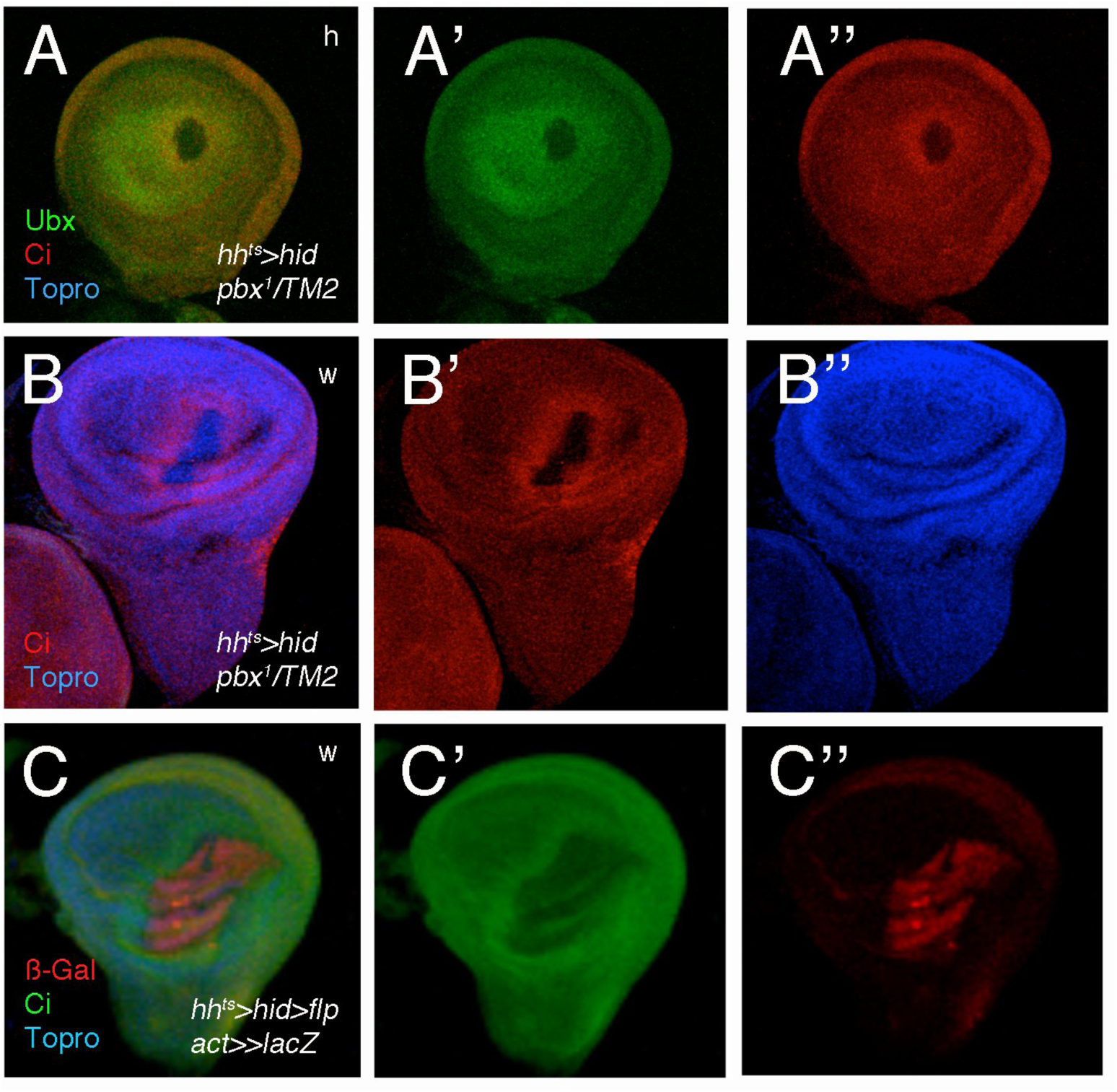
Partial duplication of anterior tissue in the wing disc after cell death induction in the posterior compartment. (A-A’’) *pbx* haltere disc in which cell death was induced in the posterior compartment (genotype UAS-*hid tub*-Gal80^ts^/+; *pbx^1^ hh*-Gal4/*TM2*). Ubx and Ci expression occupy most of the disc, with a small group of posterior cells in the middle of the pouch. (B-B’’) Wing disc of the same genotype as in A, which similarly shows anterior cells (expressing Ci) surrounding posterior cells in the middle of the pouch. (C-C’’) Wing disc in which cell death was induced in the posterior compartment and cell lineage of posterior cells are marked by ß-galactosidase signal (genotype UAS-*hid tub*-Gal80^ts^/+; *hh*-Gal4/ UAS-*flp act*>stop>*lacZ*). Similarly to the disc in B, the posterior compartment is reduced to a group of cell in the middle of the pouch and hinge, surrounded by anterior cells, expressing Ci.

Given these results, we carried out a control experiment, analyzing regeneration in wing discs without any BX-C mutation and with the same experimental conditions as in previous experiments. The genotype of the 18 discs analyzed was UAS-*hid tub*-Gal80^ts^/*+*; *hh*-Gal4/UAS-*flp act*>stop>*lacZ*. Although we found normal regeneration in most discs, some presented overlap of posterior lineage cells (ß-galactosidase-expressing cells) and Ci, or irregular A/P boundary (Fig. S6C), as previously described (Herrera and Morata, 2014). We also found discs which a reduced P compartment (Fig. S6D-D’’) and, surprisingly, one disc in which the posterior compartment is reduced and surrounded by anterior cells, a duplication similar to those observed in *pbx* wing or haltere discs (Fig. 7C-C’’) (Table 2). These results suggest that in larvae wildtype for *Ubx* function there could be, though very infrequently, partial extension of the anterior compartment and little regeneration in the posterior one.

## Discussion

In Drosophila, the existence of compartments is not an absolute barrier for regeneration (Herrera and Morata, 2014). We have tested if differences in Hox expression could result in a complete compartment/segment restriction to regeneration. Such differences have been proposed to be sufficient in some cases for the segregation of cells at *Drosophila* compartment boundaries or vertebrate rhombomeres (Curt et al., 2013; Prin et al., 20014), processes normally taken by the *engrailed*/*hedgehog*/*cubitus interruptus* system in Drosophila (Wang and Dahmann, 2020), or genes like *Kros20* in vertebrate rhombomeres (Addison and Wilkinson, 2016). Our experiments in genital and haltere discs (in a *bx* mutant) suggest that differences in Hox gene expression are not an absolute impediment for the crossing of cells to assist in regeneration, even though in these experiments the signals normally restricting this crossing are most likely still present.

We observed a different result in a *pbx* mutant background. In this case, a substantial proportion of discs present the posterior compartment reduced (sometimes quite dramatically) and the anterior cells surrounding the damaged tissue. These cases suggest that anterior cells do not contribute (or little do so) to the regeneration of the posterior compartment, and the variable regeneration from posterior cells that are not dead probably results in the different extent of posterior compartment size. This suggests that the differences in *Ubx* expression between compartments might reduce or prevent the movement of anterior cells to the posterior compartment. Surprisingly, these effects on regeneration are also observed in the wing disc, where *Ubx* is not present in the proper disc epithelium, suggesting the effect of the *pbx* mutant combination is something unrelated to *Ubx*.

Further, the effects of reduction of one compartment and its being surrounded by cells from the adjacent one are also observed, in one case, when anterior cells are undergoing cell death in a *bx* mutant background and, interestingly, in one wing disc in a larva with no *Ubx* mutation. We also noticed several examples of discs with reduced posterior compartment in *Ubx^+^* larvae, which we interpret as possible instances of limited regeneration due to a reduction in the number of anterior cells crossing the boundary. Nevertheless, the few extreme examples we have found in which the posterior compartment of haltere discs is reduced to a few cells (see Fig. 5E) cannot rule out that differences in *Ubx* expression may contribute to prevent anterior cells crossing the compartment boundary for regeneration.

Taken together, our results suggest that, occasionally, one compartment has a reduced ability to regenerate, probably because cells from the other compartment cannot readily overcome the compartment boundary, so that the damaged compartment has a reduced size. Strangely, the *pbx* mutant combination significantly increases the frequency of this effect in both haltere and wing discs, suggesting something unspecific in this mutant combination prevents a normal regeneration.

## Material and Methods

### Drosophila Stocks

The *Drosophila melanogaster* wildtype line used is Vallecas. *bx^3^, pbx^1^* and *TM2, Ubx^130^* (Lewis, 1963) are mutations in the *Ubx* gene; TRE-DsRed is a reporter of JNK activity (Chatterjee and Bohmann, 2012) and *BRV-B-GFP* is an enhancer of *wingless* that also responds to JNK activity (Harris et al., 2016); the Gal4/UAS system (Brand and Perrimon, 1993) was used with the following Gal4 lines: *cad-*Gal4^459^ (Calleja et al., 1996)*, hh-*Gal4 (Tanimoto et al., 2000), *ci*-Gal4 (Croker et al., 2006), *tsh*-Gal4 (Calleja et al., 1996), *sd*-Gal4 (Calleja et al., 1996), and the following UAS lines: UAS-*GFP* (Ito et al., 1997) (BDSC 6874), UAS-*lacZ* (Brand and Perrimon, 1993), UAS-*hid* (Igaki et al., 2000), UAS-*rpr* (Igaki et al., 2000), UAS-*y^+^* (Calleja et al., 1996) and UAS-*flp* (BDSC 4539 and BDSC 4540). The *CyO, Tb* balancer has the *Tubby* (*Tb*) dominant marker inserted on the second chromosome (Lattao et al., 2011), and was used, as the *TM6B, Tb* balancer on the third chromosome, to identify mutant larvae for disc dissection. Temporal control of gene activity in the Gal4/UAS system was achieved with *tub*-Gal80^ts^ lines (McGuire et al., 2003) (BDSC 7017 and BDSC 7019). Larvae were normally shifted from 18°C to 30°C during second or early third larval instar.

### Flip-out clones

Cell lineage clones were generated using the flip-out technique and the following stocks: *act*>*stop*>*lacZ* (Struhl and Basler, 1993), *ubi*>stop>*GFP^nls^* (BDSC32250, BDSC32251) and *act*>*y^+^*>Gal4 (Ito et al., 1997) (BDSC 3953). Clones were inducing by setting the larvae in a water bath 10 min at 37°C after 4-5 days of development at 18°C.

### Analysis of adult cuticles

Flies were dissected and macerated in a 10% KOH solution at 100°C to remove the internal structures. The cuticles were mounted in glycerol, adding a small amount of Tween-20.

### Regeneration experiments

For regeneration studies in the analia we followed this protocol: after 24h egg laying at 18°C, larvae were left growing for 5 or 6 days at 18°C, then shifted to 30°C for 30h or 40h (for experiments expressing *hid*) or 4h 30min (for experiments expressing *rpr*) and shifted back to 18°C for recovery during 2-4 days for disc dissection, or until adult eclosion. For studies in haltere and wing discs, females were laying eggs for 24h at 18°C, larvae were left growing for 6 days at 18°C, then shifted to 30°C for 30h and shifted back to 18°C for 2-4 days for disc dissection.

### Dissection and antibody staining

Larvae were dissected in cold PBS and fixation was carried out in 4% PFA for 30 min. Dissection, fixation and staining of imaginal discs were done according to standard protocols (Sullivan et al., 2000). The antibodies used were mouse anti-Ubx 1:10 (White and Wilcox, 1984), rabbit anti-ßgalactosidase 1:2000 (Cappel), mouse anti-ßgalactosidase 1:100 (Sigma), mouse anti-Abd-B 1:125 (DSHB) (Celniker et al., 1989), rat anti-Ci 1:200 (DSHB 2A1) (Motzny and Holmgreen, 1995), rabbit anti-Cad 1:200 (McDonald and Struhl, 1986), guinea-pig anti-Cad 1:200 (Kosman et al., 1988) and rabbit anti-Dcp1 1:200 (Cell Signaling, antibody #9578). Secondary antibodies were fluorescently labeled secondary antibodies (Jackson Immunoresearch) used at 1:200 dilution. DAPI (MERCK) and TO-PRO3 (Invitrogen) were used in a 1:1000 dilution to label nuclei.

### Image acquisition

Photographs of adult flies were taken with a Leica MZ12 stereomicroscope and a Leica DFC5000 camera, and images were acquired using Leica LAS software (3.7). Images from mounted adults were taken with a Zeiss Axiophot microscope. Confocal images were taken in the following microscopes: Zeiss LSM710, Zeiss LSM810, Zeiss LSM710 and Nikon A1R+ Eclipse Ti-E. Image treatment and analysis was performed using Image J and Adobe Photoshop and Powerpoint softwares.

### Statistical analysis

To compare bristle number between *cad*-Gal4 *tub*-Gal80^ts^ UAS-*flp* UAS-*GFP act*>*y^+^*>Gal4 and *cad*-Gal4 *tub*-Gal80^ts^ UAS-*flp* UAS-*hid act*>*y^+^*>Gal4, the two-tailed non-parametric Student’s *t*-test test was used, using the Graph Pad Prism software. Symbol to indicate significance was 0.01<*p<0.05.

## Acknowledgments

We thank D. Bohmann, S. Celniker, I. Hariharan, H. Lipshitz, P McDonald, G. Morata, K. Saito and R. White for stocks and antibodies. We thank Mar Casado and Nuria Esteban for stocks care and curation and the Confocal Microscopy Service at CBMSO for help in the analysis of images. We also thank the Bloomington Stock Center, the Asian Distribution Center for Segmentation Antibodies-Invertebrate Genetics Laboratory, Shizuoka, Japan, and the Developmental Studies Hybridoma Bank for fly stocks and reagents. This study was supported by grants from FEDER/Ministerio de Ciencia e Innovación-Agencia Estatal de Investigación-Consejo Superior de Investigaciones Científicas (No. BFU2014-51989-P, BFU2017-86244-P and PID2020-113318GB-I00). R.A. J. was a recipient of a CONACyT Fellowship from the Mexican Academy of Sciences (MAS) and the National Council for Science and Technology (CONACYT). Institutional support from the Ramón Areces Foundation is acknowledged.

## References

Addison M., Wilkinson D. G. (2016). Segment Identity and Cell Segregation in the Vertebrate Hindbrain. Current Topics in Developmental Biology 117:581–96. doi: 10.1016/bs.ctdb.2015.10.019.

Aza-Blanc P., Ramírez-Weber F. A., Laget M. P., Schwartz C., Kornberg T. B. (1997). Proteolysis that is inhibited by hedgehog targets Cubitus interruptus protein to the nucleus and converts it to a repressor. Cell 89: 1043–53. doi: 10.1016/s0092-8674(00)80292-5.

Beachy P. A., Helfand S. L., Hogness, D. S. (1985). Segmental distribution of bithorax complex proteins during Drosophila development. Nature 313: 545–551. doi: 10.1038/313545a0.

Bergantinos C., Corominas M., Serras F. (2010). Cell death-induced regeneration in wing imaginal discs requires JNK signalling. Development 137: 1169–1179. doi: 10.1242/dev.045559.

Bosch J. A., Sumabat T. M., Hariharan I. K. (2016). Persistence of RNAi-Mediated Knockdown in Drosophila Complicates Mosaic Analysis Yet Enables Highly Sensitive Lineage Tracing. Genetics 203: 109–18. doi: 10.1534/genetics.116.187062.

Brand A. H., Perrimon N. (1993). Targeted gene expression as a means of altering cell fates and generating dominant phenotypes. Development 118: 401–15. doi: 10.1242/dev.118.2.401.

Bryant P. J. (1975a). Regeneration and duplication in imaginal discs. Ciba Foundation Symposium 29: 71–93. doi: 10.1002/9780470720110.ch5.

Bryant P. J. (1975b). Pattern formation in the imaginal wing disc of Drosophila melanogaster: Fate map, regeneration and duplication. Journal of Experimental Zoology 193: 49–77. doi: 10.1002/jez.1401930106.

Cabrera C. V., Botas J., Garcia-Bellido A. (1985). Distribution of Ultrabithorax proteins in mutants of Drosophila bithorax complex and its transregulatory genes. Nature 318: 569–571. doi10.1038/328569a0.

Calleja M., Moreno E., Pelaz S., Morata G. (1996). Visualization of gene expression in living adult Drosophila. Science 274: 252–5. doi: 10.1126/science.274.5285.252.

Casares F., Sánchez L., Guerrero I., Sánchez-Herrero E. (1997). The genital disc of Drosophila melanogaster. : I. Segmental and compartmental organization. Development Genes and Evolution. 207: 216–228. doi: 10.1007/s004270050110.

Celniker S. E., Keelan D. J., Lewis E. B. (1989). The molecular genetics of the bithorax complex of Drosophila: characterization of the products of the Abdominal-B domain. Genes & Development 3: 1424–36. doi: 10.1101/gad.3.9.1424.

Chatterjee N., & Bohmann D. (2012). A Versatile ΦC31 Based Reporter System for Measuring AP-1 and Nrf2 Signaling in Drosophila and in Tissue Culture. PLoS ONE 7: e34063. doi: 10.1371/journal.pone.0034063.

Croker J. A., Ziegenhorn S. L., Holmgren, R. A. (2006). Regulation of the Drosophila transcription factor, Cubitus interruptus, by two conserved domains. Developmental Biology 291: 368–81. doi: 10.1016/j.ydbio.2005.12.020.

Curt J. R., de Navas L. F., Sánchez-Herrero E. (2013). Differential activity of Drosophila Hox genes induces myosin expression and can maintain compartment boundaries. PLoS One 8: e57159. doi: 10.1371/journal.pone.0057159.

Dübendorfer K., Nöthiger R. (1982). A clonal analysis of cell lineage and growth in the male and female genital disc of Drosophila melanogaster. Wilhelm Roux’s Archives of Developmental Biology 191: 42–55. doi: 10.1007/BF00848545.

Eaton S., Kornberg T. B. (1990). Repression of ci-D in posterior compartments of Drosophila by engrailed. Genes & Development 6: 1068–77. doi: 10.1101/gad.4.6.1068.

Estrada B., Sánchez-Herrero E. (2001). The Hox gene Abdominal-B antagonizes appendage development in the genital disc of Drosophila. Development 128: 331–339. doi: 10.1242/dev.128.3.331.

Foronda D., Estrada B., de Navas L., Sánchez-Herrero E. (2006). Requirement of abdominal-A and Abdominal-B in the developing genitalia of Drosophila breaks the posterior downregulation rule. Development 133: 117–127. doi: 10.1242/dev.02173.

Freeland D. E., Kuhn D. T. (1996). Expression patterns of developmental genes reveal segment and parasegment organization of D. melanogaster genital discs. Mechanisms of Development 56: 61–72. doi: 10.1016/0925-4773(96)00511-4.

Gandille P., Narbonne-Reveau K., Boissonneau E., Randsholt N., Busson D., Pret A. M. (2010). Mutations in the polycomb group gene polyhomeotic lead to epithelial instability in both the ovary and wing imaginal disc in Drosophila. PLoS One 5:e13946. doi: 10.1371/journal.pone.0013946.

Garcia-Bellido A. (1968). Cell affinities in antennal homoeotic mutants of Drosophila melanogaster. Genetics 59: 487–99. doi: 10.1093/genetics/59.4.487.

Garcia-Bellido A., Ripoll P., Morata G. (1973). Developmental compartmentalisation of the wing disk of Drosophila. Nature New Biology 245: 251–3. doi: 10.1038/newbio245251a0.

Garg A., Srivastava A., Davis M. M., O’Keefe S. L., Chow L., Bell J. B. (2007). Antagonizing Scalloped with a Novel Vestigial Construct Reveals an Important Role for Scalloped in *Drosophila melanogaster* Leg, Eye and Optic Lobe Development. Genetics 175: 659–69. doi: 10.1534/genetics.106.063966.

Gorfinkiel N., Sánchez L., Guerrero I. (2003). Development of the Drosophila genital disc requires interactions between its segmental primordia. Development (Cambridge, England), 130: 295–305. doi: 10.1007/BF01938454.

Harris R. E., Setiawan L., Saul J., Hariharan I. K. (2016). Localized epigenetic silencing of a damage-activated WNT enhancer limits regeneration in mature Drosophila imaginal discs. Elife 5: e11588. doi: 10.7554/eLife.11588.

Herrera S. C., Martín R., Morata G. (2013). Tissue homeostasis in the wing disc of Drosophila melanogaster: immediate response to massive damage during development. PLoS Genetics 9: e1003446. doi: 10.1371/journal.pgen.1003446.

Herrera S. C., Morata G. (2014). Transgressions of compartment boundaries and cell reprogramming during regeneration in Drosophila. eLife 3: e01831. doi: 10.7554/eLife.01831.

Igaki T., Kanuka H., Inohara N., Sawamoto K., Núñez G., Okano H., Miura M. (2000). Drob-1, a Drosophila member of the Bcl-2/CED-9 family that promotes cell death. Proceedings of the National Academy of Sciences of the United States of America 97: 662–7. doi: 10.1073/pnas.97.2.662.

Ito K., Awano W., Suzuki K., Hiromi Y., Yamamoto D. (1997). The Drosophila mushroom body is a quadruple structure of clonal units each of which contains a virtually identical set of neurones and glial cells. Development 124: 761–71. doi: 10.1242/dev.124.4.761.

Kosman D., Small S., Reinitz J. (1998). Rapid preparation of a panel of polyclonal antibodies to Drosophila segmentation proteins. Development Genes and Evolution 208: 290–4. doi: 10.1007/s004270050184.

Lattao R., Bonaccorsi S., Guan X., Wasserman S. A., Gatti M. (2010). Tubby-tagged balancers for the Drosophila X and second chromosomes. Fly 5: 369–70. doi: 10.4161/fly.5.4.17283.

Lewis E. B. (1963). Genes and Developmental Pathways. American Zoologist 3: 33–56. https://doi.org/10.2307/3881152

Lewis E. B. (1978). A gene complex controlling segmentation in Drosophila. Nature 276: 565–70. doi: 10.1038/276565a0.

Littlefield C. L., Bryant P. J. (1979). Prospective fates and regulative capacities of fragments of the female genital disc of *Drosophila melanogaster*. Developmental Biology 70: 127–148. doi: 10.1016/0012-1606(79)90012-5.

Macías A., Romero N. M., Martín F., Suárez L., Rosa A. L., Morata G. (2004). PVF1/PVR signaling and apoptosis promotes the rotation and dorsal closure of the Drosophila male terminalia. International Journal of Developmental Biology 48: 1087–94. doi: 10.1387/ijdb.041859am.

Macdonald P. M., Struhl G. (1986). A molecular gradient in early Drosophila embryos and its role in specifying the body pattern. Nature 324: 537–45. doi: 10.1038/324537a0.

McGuire S. E., Le P. T., Osborn A. J., Matsumoto K., Davis, R. L. (2003). Spatiotemporal Rescue of Memory Dysfunction in Drosophila. Science 302: 1765–1768. doi10.1126/science.1089035.

McQueen E., Rice G., Pillai S., Ziabari O. S., Vincent B. J., Rebeiz M. (2026). Parallels in the Regulatory Landscape of Dimorphic Female and Male Genital Structures in *Drosophila melanogaster*. Genetics 232: iyaf221. doi:10.1093/genetics/iyaf221.

Mlodzik M., Gehring W. J. (1987). Expression of the caudal gene in the germ line of Drosophila: formation of an RNA and protein gradient during early embryogenesis. Cell 48: 465–78. doi: 10.1016/0092-8674(87)90197-8.

Morata G., Garcia-Bellido A. (1976). Developmental analysis of some mutants of the bithorax system of Drosophila. Wilhelm Roux’s Archives of Developmental Biology 179: 125–143. doi: 10.1007/BF00848298.

Moreno E., Morata G. (1999). Caudal is the Hox gene that specifies the most posterior Drosophile segment. Nature 400: 873–877. doi10.1038/23709.

Motzny C. K., Holmgreen R. (1995). The Drosophila cubitus interruptus protein and its role in the wingless and hedgehog signal transduction pathways. Mechanisms of Development 52: 137–50. doi: 10.1016/0925-4773(95)00397-j.

Nöthiger R., Dübendorfer A., Epper F. (1977). Gynandromorphs reveal two separate primordia for male and female genitalia inDrosophila melanogaster. Wilhelm Roux’s Archives of Developmental Biology 181: 367–373. doi10.1007/BF00848062

Paul R, Giraud G, Domsch K, Duffraisse M, Marmigère F, Khan S, Vanderperre S, Lohmann I, Stoks R, Shashidhara LS, Merabet S. (2021). Hox dosage contributes to flight appendage morphology in Drosophila. Nature Communications 12: 2892. doi: 10.1038/s41467-021-23293-8.

Prin F., Serpente P, Itasaki N, Gould AP. (2014). Hox proteins drive cell segregation and non-autonomous apical remodelling during hindbrain segmentation. Development 141: 1492–502. doi: 10.1242/dev.098954.

Repiso A., Bergantiños C., Serras F. (2013). Cell fate respecification and cell division orientation drive intercalary regeneration in Drosophila wing discs Development. 140: 3541–51. doi: 10.1242/dev.095760.

Rezsohazy R., Saurin A. J., Maurel-Zaffran C., Graba Y. (2015). Cellular and molecular insights into Hox protein action. Development 142: 1212–27. doi: 10.1242/dev.109785.

Rousset R., Bono-Lauriol S., Gettings M., Suzanne M., Spéder P., Noselli S. (2010). The Drosophila serine protease homologue Scarface regulates JNK signalling in a negative-feedback loop during epithelial morphogenesis. Development 137: 2177–86. doi: 10.1242/dev.050781.

Schüpbach T., Wieschaus E., Nöthiger R. (1978). The embryonic organization of the genital disc studied in genetic mosaics of *Drosophila melanogaster*. Wilhelm Roux’s Archives of Developmental Biology 185: 249–270. doi10.1007/BF00848355

Smith-Bolton R. K., Worley M. I., Kanda H., Hariharan I. K. (2009). Regenerative Growth in Drosophila Imaginal Discs Is Regulated by Wingless and Myc. Developmental Cell 16: 797–809. doi: 10.1016/j.devcel.2009.04.015.

Struhl G. Basler K. (1993). Organizing activity of wingless protein in Drosophila. Cell 72: 527–40. doi: 10.1016/0092-8674(93)90072-x.

Sullivan W., Ashburner M. Hawley R. S. (2000). *Drosophila* Protocols. Cold Spring Harbor Laboratory Press, New York.

Tanimoto H., Itoh S., ten Dijke P., Tabata T. (2000). Hedgehog creates a gradient of DPP activity in Drosophila wing imaginal discs. Molecular Cell 5: 59–71. doi: 10.1016/s1097-2765(00)80403-7.

Tiberghien M. A., Lebreton G., Cribbs D., Benassayag C., Suzanne M. (2015). The Hox gene Dfd controls organogenesis by shaping territorial border through regulation of basal DE-Cadherin distribution, Developmental Biology 405: 183–8. doi: 10.1016/j.ydbio.2015.07.020.

Ursprung H. (1962). Influence of host age on the extent of development of sagitally fragmented male genital disks of phila melanogaster. Developmental Biology 4: 22–39. doi: 10.1016/0012-1606(62)90031-3.

Verghese S., Su T. T. (2016). Drosophila Wnt and STAT Define Apoptosis-Resistant Epithelial Cells for Tissue Regeneration after Irradiation. PLoS Biology 14: e1002536. doi: 10.1371/journal.pbio.1002536.

Wang J., Dahmann C. (2020). Establishing compartment boundaries in Drosophila wing imaginal discs: An interplay between selector genes, signaling pathways and cell mechanics Seminars in Cell and Developmental Biology107: 161-169. doi: 10.1016/j.semcdb.2020.07.008.

White R. A., Wilcox M. (1984). Protein products of the bithorax complex in Drosophila. Cell 39: 163–171. doi: 10.1016/0092-8674(84)90202-2.

White R. A., Wilcox M. (1985). Distribution of Ultrabithorax proteins in Drosophila. EMBO Journal 8: 2035–43. doi: 10.1002/j.1460-2075.1985.tb03889.x.

Worley M. I., Setiawan L., & Hariharan I. K. (2012). Regeneration and Transdetermination in Drosophila Imaginal Discs. Annual Review of Genetics 46: 289–310. 10.1146/annurev-genet-110711-155637.

