## Supplemental Figures for "Regeneration can take place across Drosophila compartments or segments with different Hox gene expression"

### Supplementary Figures

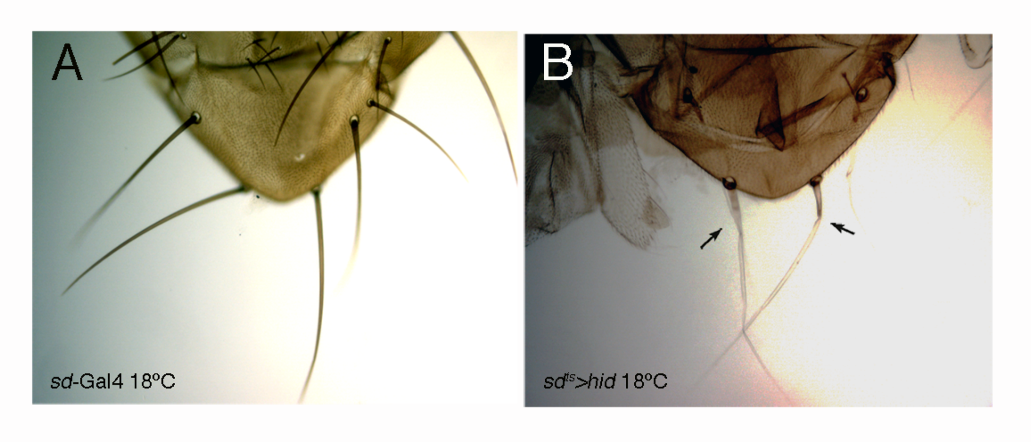


**Supplementary Figure 1. Effect of *hid* expression with *tub*-Gal80$^{ts}$ on bristle development at 18ºC.** (A) scutellum of a wildtype fly. (B) Scutellum of a *scalloped*-Gal4; UAS-*hid* tub-Gal80$^{ts}$/+ fly grown at 18ºC, showing thin and near transparent bristles (*scalloped* is expressed in bristles, Garg et al., 2007).

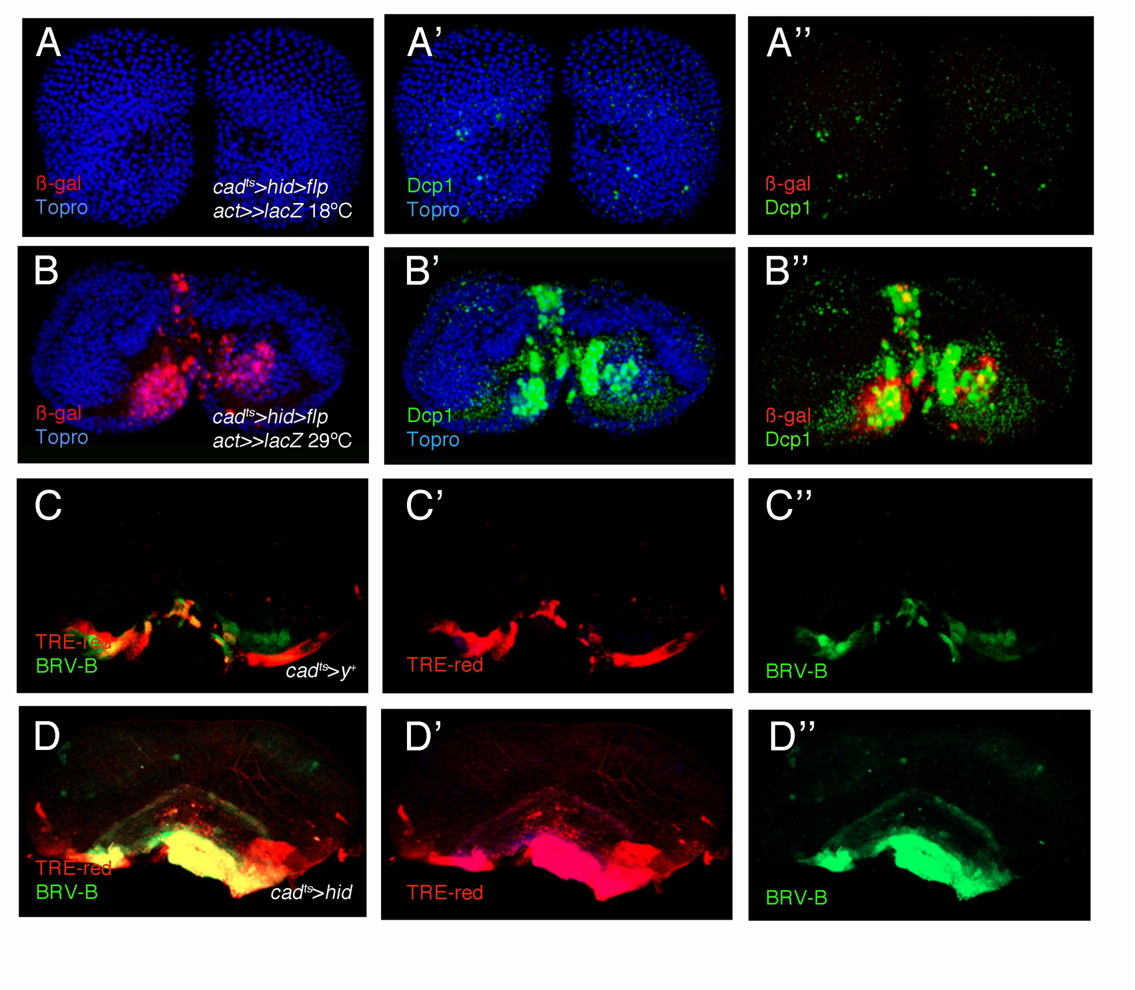

**Supplementary Figure 2. Caspase, JNK pathway and wingless reporter activation after induction of cell death in the genital disc.** (A-B'') Cleaved Dcp1 staining (in green) and ß-galactosidase signal (in red) in the distorted A10 of third instar larvae male genital discs of genotype *cad*-Gal4/UAS-*hid tub*-Gal80$^{ts}$; UAS-*flp act*>stop>*lacZ*/+, in which *cad* lineage expression and apoptosis are either not induced (control discs, A-A'') because the larvae are grown at 18ºC, or induced (B-B''), when they are shifted to 30ºC. Topro is in blue. (C-C'') *cad*-Gal4 *tub*-Gal80$^{ts}$/UAS-*y*$^{+}$; BRV-B-GFP TRE-DsRed/+ third instar male genital disc (larvae shifted from 18ºC to 30ºC), in which JNK activation (TRE-DsRed reporter) or BRV-B-GFP wg-reporter signal (in green) is limited to the stalk region of the disc. (D-D'') In *cad*-Gal4 *tub*-Gal80$^{ts}$/UAS-*hid*; BRV-B-GFP TRE-DsRed/+ third instar male genital disc (larvae shifted from 18ºC to 30ºC) both TRE-DsRed and wg reporter expression is extended in the A10 segment.

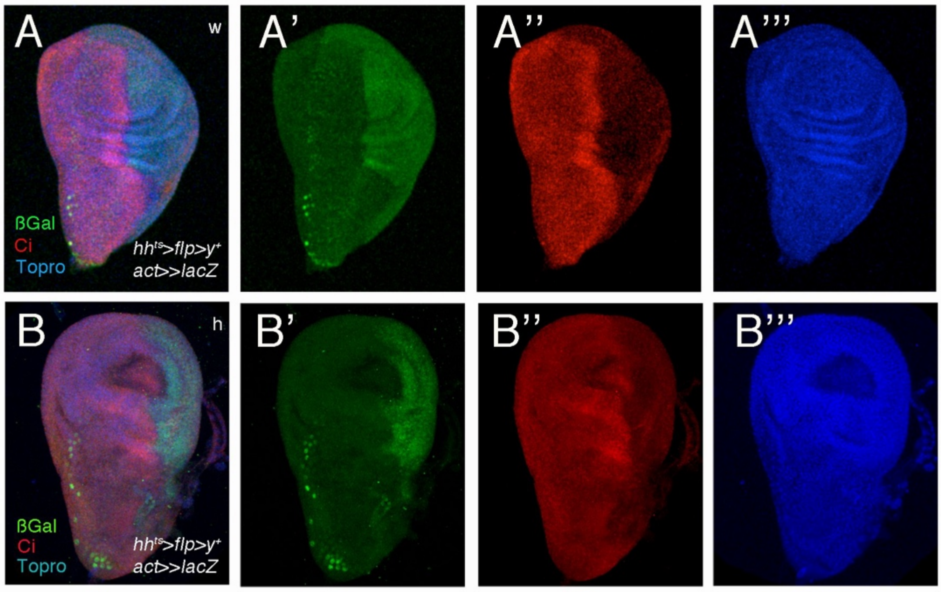


**Supplementary Figure 3. Complementary expression of *cubitus interruptus* and posterior compartment cell lineage**. (A-B''') Wing (w) (A-A''') and haltere (h) (B-B''') third instar discs showing Ci expression in the anterior compartment complementary to the cell lineage of the posterior compartment, marked by expressing lacZ through cassette recombination (genotype: UAS-*y*$^{+}$/+; UAS-*flp act*>

*stop*>*lacZ*/*hh*-Gal4 *tub*-Gal80$^{ts}$). Second instar larvae were shifted from 18ºC to 30ºC. Topro is in blue.

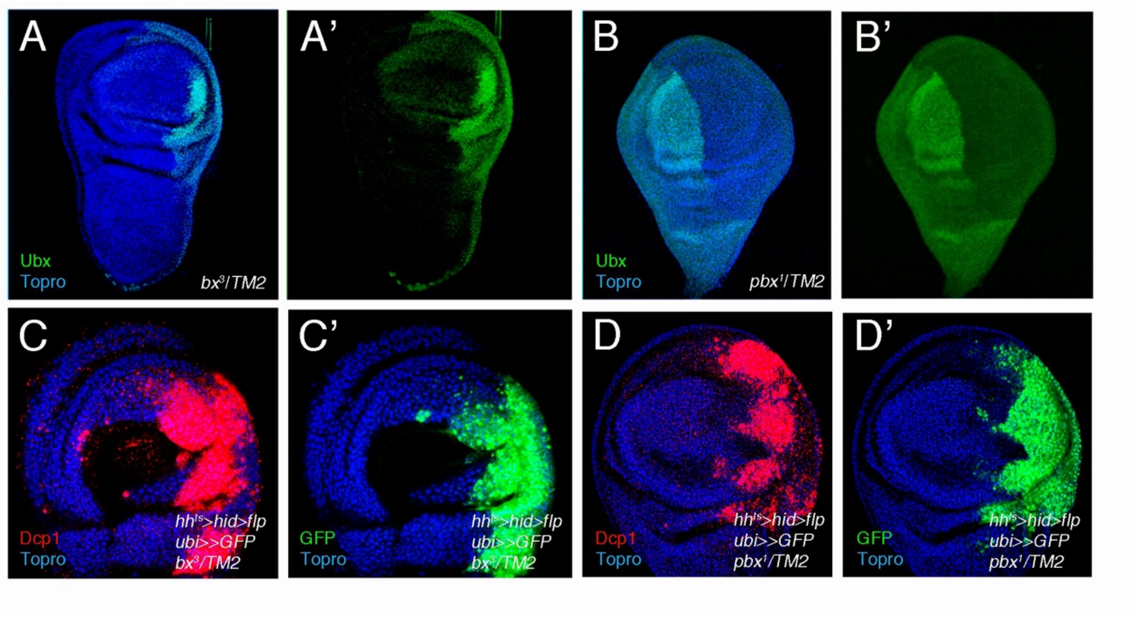


**Supplementary Figure 4. *Ubx* expression and cell death induction in *bx* and *pbx* mutants.** (A-B'). *Ubx* expression in *bx*$^3$/*TM2* (A, A') and *pbx*$^1$/*TM2* (B, B') haltere discs. Topro is in blue. (C-D') Dcp1 signal (C, D) and cell lineage expression (C', D') after cell death induction with *hid* in the posterior compartment of *bx*$^3$/*TM2* (C, C') and *pbx1/TM2* (D, D') haltere discs (*ubi*>stop>*GFP* UAS-*flp*/UAS-*hid tub*-Gal80$^{ts}$; *bx*$^3$ or *pbx*$^1$ *hh*-Gal4/*TM2*). Treatment of discs in panel C was 48h 30ºC, and for D it was

30h 30ºC.

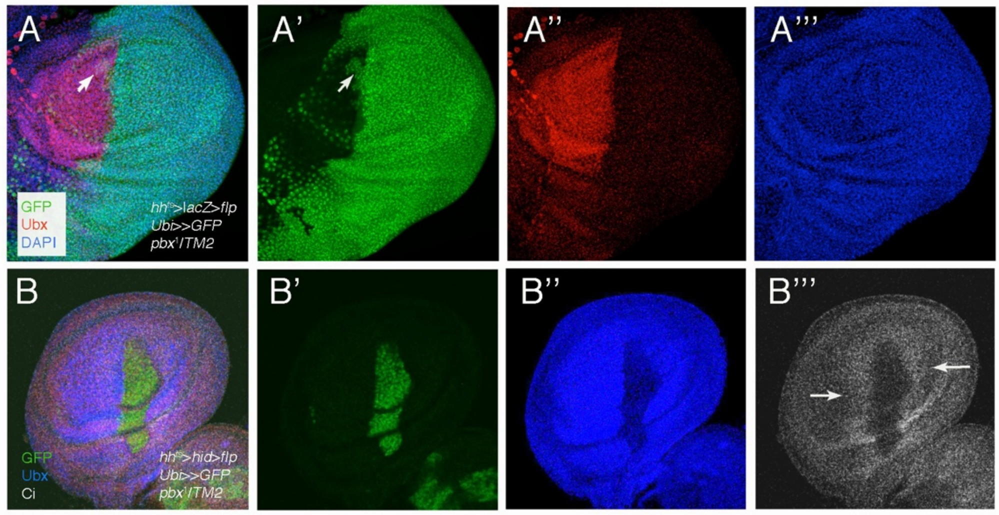


**Supplementary Figure 5. Cell lineage and *Ubx* expression in control and experimental (*hid* expression) *pbx* mutant discs.** (A-A’’’) Cell lineage and Ubx antibody signal after expression of *lacZ* in the posterior compartment of *pbx*$^{1}$/*TM2* haltere discs (*ubi*>stop>*GFP* UAS-*flp*/UAS-*lacZ tub*-Gal80$^{ts}$; *bx*$^{3}$ *hh*-Gal4/*TM2*). Note a group of posterior cells overlapping Ubx signal (arrow). (B-B’’’) Cell lineage and Ubx and Ci antibody signals after induction of cell death in the posterior compartment of *pbx*$^{1}$/*TM2* haltere discs (*ubi*>stop>*GFP* UAS-*flp*/UAS-*hid tub*-Gal80$^{ts}$; *pbx*$^{1}$ *hh*-Gal4/*TM2*). The posterior cells are surrounded by anterior ones.

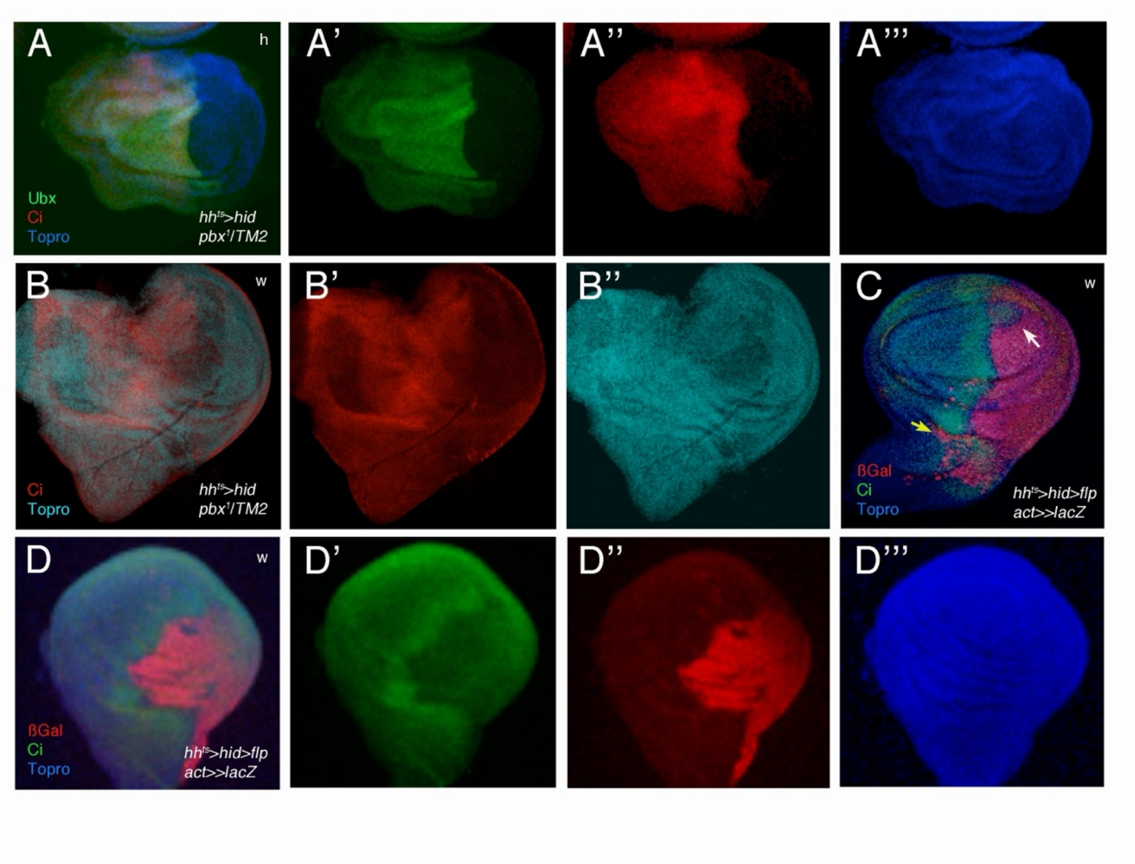


**Supplementary Figure 6. Abnormal development of wing and haltere discs after induction of cell death in the posterior compartment.** (A-B') Haltere (A-A''') or wing (B-B'') *pbx*$^1$/*TM2* discs in which *hid* was induced in the posterior compartment, stained with Ubx and Ci (UAS-*hid tub*-Gal80$^{ts}$/+; *hh*-Gal4 *pbx*$^1$/*TM2*). (C) Wing disc in which cell death was induced in the posterior compartment (UAS-*hid tub*-Gal80$^{ts}$/+; *hh*-Gal4/UAS-*flp act*>stop>*lacZ*). The cell lineage of posterior cells is detected by ß-galactosidase expression and anterior cells by Ci signal. Note anterior cells entering the posterior compartment (white arrow) and posterior cells entering the anterior one (yellow arrow). (D-D''') Wing disc in which cell death was induced in the posterior compartment (UAS-*hid tub*-Gal80$^{ts}$/+; *hh*-Gal4/UAS-*flp act*>stop>*lacZ*). The cell lineage of posterior cells is detected by ß-galactosidase expression and anterior cells by Ci signal. The posterior compartment is highly reduced and the antero-posterior boundary is irregular.
